# Dual expansion routes likely underlie the present-day population structure in a *Parnassius* butterfly across the Japanese Archipelago

**DOI:** 10.1101/2024.07.16.603823

**Authors:** Hideyuki Tamura, Tomoaki Noda, Mikiko Hayashi, Yuko Fujii, Noriko Iwata, Yuko Yokota, Masanori Murata, Chisato Tatematsu, Hideshi Naka, Akio Tera, Katsumi Ono, Kakeru Yokoi, Takanori Kato, Tomoko Okamoto, Koji Tsuchida

## Abstract

The Japanese Archipelago, comprising a series of isolated yet interconnected islands, had been geographically separated from the Eurasian continent. The linear topography presents a unique biogeographic context for dispersing organisms from the continent. In this study, we utilized mitochondrial DNA (mtDNA) and single nucleotide polymorphism (SNP) variation to elucidate the dispersal history of the Japanese clouded butterfly *Parnassius glacialis* across the Japanese Archipelago, including North Island (Hokkaido), Main Island (Honshu) and Shikoku Island. Our analysis of mtDNA (COI, COII) at 1192 base pairs (bps) revealed 49 haplotypes and identified three distinct haplotype groups in the network. These groups correspond geographically to East Japan, West Japan, and Chugoku-Shikoku. The Chugoku-Shikoku group is the most ancient lineage. Interestingly, the Chugoku-Shikoku lineage showed a closer network connection to the East Japan lineage than the geographically proximate West Japan lineage. Divergence time estimates suggest that the Chugoku-Shikoku lineage diverged from the continental *P. glacialis* approximately 3.05 million years ago (MYA). Subsequently, from the Chugoku-Shikoku lineage, the East Japan and West Japan lineages diverged around 1.05 MYA, with the subsequent divergence between the East and West Japan lineages occurring at approximately 0.62 MYA. Based on the 3067 SNP genotypes, population structure analysis revealed five distinct genetic structures within the Japanese Archipelago, indicating geographical differentiation. From the analyses by mtDNA and SNP variations, four primary genetic barriers were identified: between Hokkaido and Honshu, between East and Central Japan, within the Kansai region, and within the Chugoku region. The former three lines corresponded to the Blakiston Line, the Itoigawa-Shizuoka Tectonic Line, and Lake Biwa, respectively. These findings suggest that *P. glacialis* diverged from the continental *P. glacialis* and expanded its range across the Japanese Archipelago via the North and South routes, establishing its current distribution.

## 1. Introduction

Evolutionary biological factors, including distances between individuals and populations, stochastic events, and local population adaptations (e.g., Wright, 1943; Nosil and Crespi, 2004; Nosil et al., 2005), reflect relatively recent events. On the other hand, geological factors encompassing the formation of historical landforms reflect considerably older events than the former (e.g., Otofuji et al., 1985; Jolivet et al., 1994; Martin, 2011). The distribution of organisms is influenced by these two main factors, resulting in their current distribution patterns. Indeed, organisms carry the history of these influences.

The Proto-Japanese archipelago islands were located on the eastern edge of the Eurasian continent until 20 million years ago (MYA), constituting the easternmost part of the Eurasian continent. They originated from an accretionary wedge created by the subduction of the oceanic plate beneath the continental plate (Nakajima, 2018). According to the double-door hypothesis (Otofuji et al., 1985; Jolivet et al., 1994; Martin, 2011), two half-arcs in the south-west and the north-east were later differentiated and migrated from the continent, forming the Sea of Japan with them. These half-arcs underwent opposing rotations and moved to their approximate positions, separated by a valley known as Fossa Magna. This valley was later filled through alluvial and volcanic activity at approximately 5 MYA, connecting the two half-arcs by land and roughly forming the present-day outline of the Japanese Archipelago.

As the Japanese Archipelago is linearly connected from subarctic to subtropical zones, species expansions into the Archipelago can be divided into two main routes (Fig. 1). The first is from the West, which can be further divided into two sub-routes: via the Korean Peninsula and the Ryukyu Archipelago. A land bridge connected eastern China and western Japan continuously from the Late Miocene to the end of the Pliocene (10–1.7 MYA; million years ago), allowing continuous immigration of continental species into the Archipelago. Kitamura and Kimoto (2004) estimated the presence of a land bridge between Kyushu and the continent at least 3.1 and 1.7 MYA, followed by subsequent submergence.

**Fig 1.**
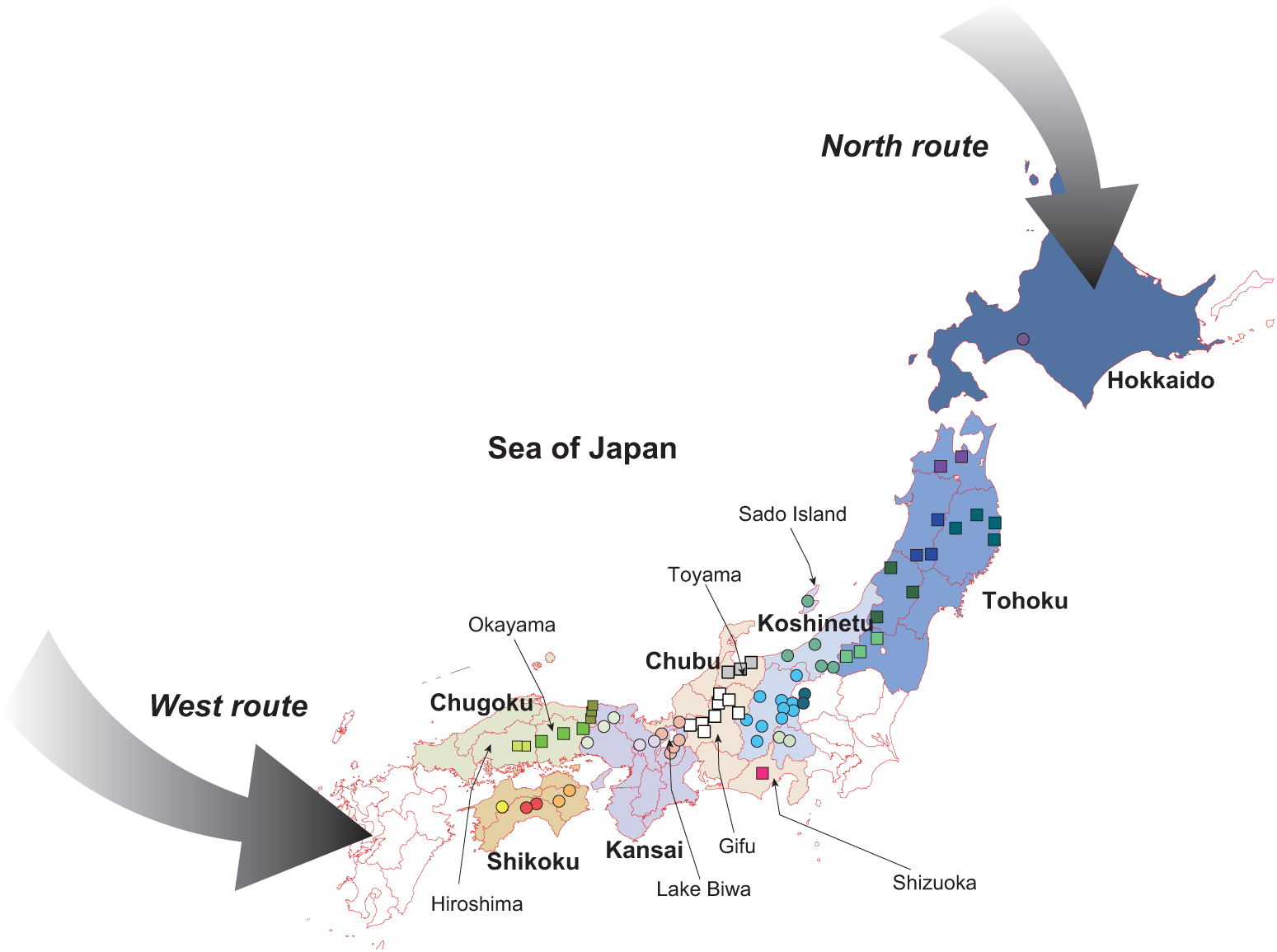
The outline of the present-day Japanese Archipelago and expected biological expansion routes from the Eurasian continent, North and West routes. The regional names are shown in bold, and prefecture names cited in the text were also shown. Circles and squares indicate each collection site for SNP analyses in this study. Samples for mtDNA were collected in Fukui and Tochigi prefectures in addition to these sites.

In contrast, the North route is also divided into two sub-routes: one via Sakhalin and the other via the Kuril Islands. The northern island of Hokkaido was repeatedly connected to the continent during the Pleistocene, including the last glacial periods (0.01−0.07 MYA, Fujimaki, 1994; Goto, 1994), with a land bridge between Sakhalin and Hokkaido facilitating the immigration of many faunal species from the continent. This land bridge is supported by evidence that the same phylogenetic insects are distributed in both Hokkaido and Sakhalin (e.g., Sota and Hayashi, 2007; Hayashi and Sota, 2014). Both the West and North routes are connected to the Eurasian continent, and it is thought that part of the lineages differentiated on the Eurasian continent spread and settled in the Japanese Archipelago. Occasionally, these lineages are dispersed from the Japanese Archipelago back to the continent via these routes (Tojo et al., 2017).

Species widely distributed across the Japanese Archipelago are excellent subjects for elucidating the processes of their spread and exploring the mechanisms underlying their local adaptation. Among these species, butterflies of the genus *Parnassius* have attracted significant interest from researchers investigating their dispersal from the continent and their migration routes. This genus originated on the Tibetan Plateau and was initially adapted to alpine environments. It is believed that *P. glacialis* differentiated as it adapted to lowland climates after experiencing a population bottleneck in the lowlands of the continent (Si et al., 2020; Su et al., 2020; Tao et al., 2020; Zhao et al., 2022, 2023; Tian et al., 2023).

*Parnassius glacialis* is a butterfly species distributed across Hokkaido (Northern Island), Honshu (Main Island), and Shikoku Island in the Japanese Archipelago. It was initially described as *P. glacialis* but has recently been referred as a separate species, *P. citrinarius* without any supporting evidence (see supplementary materials for the species name used in this study). Yagi et al. (2001) reported that populations of both *P. citrinarius* (= *glacialis*) and *P. stubbendorfii* in Japan may have been isolated from the continent on the Japanese islands during the early Pleistocene glacial period (ca. 1.7-2.0 Ma). They noted that the genetic distance between the Japanese *P. citrinarius* (= *glacialis*) and continental *P. glacialis* is substantial enough to classify them as separate species compared to the genetic distances observed between some other *Parnassius* species. Recently, Nagata (2024) also found that *P. citrinarius* (= *glacialis*) shows differentiation between eastern and western Japan within the Archipelago. However, the relatively small numbers of nucleotide sequences used in the studies have limited the ability to further clarify the detailed degree of differentiation and the expansion processes into the Japanese Archipelago.

This study aimed to analyze the DNA variations of *P. glacialis* in Japan to estimate the dispersal process from the continent and the degree of variation within the Japanese Archipelago. For this purpose, the COI and COII regions in the mitochondrial DNA (mtDNA) were analyzed. These mtDNA analyses revealed that the Japanese lineage can be divided into three main lineages. The relationship between these lineages and the continental *P. glacialis* was also examined, and the divergence dates were estimated using whole mtDNA genome sequencing. Furthermore, the recently developed GRAS-Di method (Enoki and Takeuchi, 2018; Hosoya et al., 2019) was employed to detect SNP loci in the species and to explore detailed population structure based on these loci.

## 2. Materials and Methods

### 2.1. Samples

We collected the butterflies with an insect net and taken back to the laboratory. Adult butterflies were collected at one to nine sites per prefecture, and the prefectural units were treated as a single population in this study. Their wings were excised and dried, while the remaining parts were stored in 100% ethanol (EtOH) at 4°C until DNA extraction. All samples were collected from areas where this species is not banned from collecting. From the samples, we selected 160 samples for 22 populations for SNP analyses and 185 samples for 24 populations for mtDNA analyses. Samples for mtDNA were collected in Fukui and Tochigi prefectures in addition to these sites for SNP analyses. DNA was extracted from each sample as described below.

### 2.2. Sequencing of mtDNA COI and COII

DNA was extracted from two or three legs per specimen using a NucleoSpin® Tissue kit (MACHEREY-NAGEL Düren, Germany). The isolated DNA was resuspended in Tris-HCl buffer (pH 7.5) and stored at 4 °C. Polymerase chain reaction (PCR) was conducted to amplify two gene regions: the contiguous mitochondrial cytochrome oxidase I (COI) region of 721 base pairs (bps) and cytochrome oxidase II (COII) region of 470 bps. Primers to amplify the COI region were JNfwd (5’-GCAGGAACTGGATGAACAG-3’) and Krev (5’-GAGTATCGTCGAGGTATCC-3 ’) (Dechaine et al., 2004). Primers used to amplify the COII region were PierreIII (5’-AGAGAGTTTCACCTTTAATAGAACA-3’; Nazari et al., 2007) and Eva (5’-GAGAACATTACTTGCTTTCAGTCATCT-3’; Bogdanowicz et al., 1993). Both fragments were amplified using TaKaRa Ex Taq Hot Start Version (TaKaRa, Tokyo, Japan). PCR conditions for the COI region were 94°C for 5 min, followed by 30 cycles of 94°C for 30 s, 50°C for 30 s, and 72°C for 1 min. For the COII region, 95°C for 5 min, followed by 35 cycles of 94°C for 1 min, 50°C for 1 min, 72°C for 1 min, and finally 72°C for 10 min. Sequencing reactions were performed using BigDye terminator v3.1 (Applied Biosystems, Carlsbad, USA). All samples were sequenced using an ABI PRISM 3100-Avant Genetic Analyzer (Applied Biosystems, Carlsbad, CA, USA). Sequencing was conducted at the Life Science Experimental Center and Genome Research Department, Gifu University. PCR fragments were sequenced in both directions to ensure accuracy. The sequences were aligned the sequences using MEGA 6.06 (Tamura et al., 2013).

We estimated population demography using a Bayesian skyline plot by BEAST v.2.7.6 (Drummond et al., 2006; Drummond & Rambaut, 2007). Nucleotide diversity, haplotype diversity, Fu’s F, and Tajima’s D were estimated by Arlequine 3.5.2.2 (Excoffier and Lischer 2010).

### 2.3. Whole Genome Sequencing of mtDNA for Three Lineages

Our COI and COII analyses revealed that *P. glacialis* across the Japanese Archipelago is broadly divided into the East Japan, West, and Chugoku-Shikoku lineages (see Results). Therefore, three representative individuals were selected from these three lineages (the East Japan from Hokkaido, West Japan from Shiga, and Chugoku-Shikoku from Kochi), and DNA was extracted from the thoraxes of these specimens. Extraction was carried out using the previously mentioned kit, following the manufacturer’s protocol. After DNA extraction, the following analyses were conducted at a commercial sequencing facility (Bioengineering Lab Co., Ltd., Kanagawa, Japan). Libraries were prepared according to the manual using the MGIEasy FS DNA Library Prep Set (MGI Tech). Following the provided manual, Cyclized DNA was prepared from the library and the MGIEasy Circularisation Kit (MGI Tech). DNBs were prepared according to the manual using the DNBSEQ-G400RS High-throughput Sequencing Kit (MGI Tech). Sequencing analysis of the produced DNBs was performed using the primer supplied with the High-throughput Pair-End Sequencing Primer Kit (App-D) (MGI Tech) and DNBSEQ-G400 (MGI Tech) platform at 2×150 bp. After removing adapter sequences using Cutadapt v. 4.0 (Martin, 2011), Sickle v. 1.33 (Joshi and Fass, 2011) was used to remove bases with a quality score of less than 20 and paired reads with less than 75 bases. SPAdes v. 3.15.5 (Bankevich et al. 2012) was used to assemble high-quality reads with the following parameters: a lower limit of read coverage (--cov-cutoff = 10) and mismatch correction (–careful). Mitochondrial sequences were extracted from the assembled sequences using Metaxa v. 2.2 (Bengtsson-Palme et al., 2015).

Adapter trimming and low-quality read removal were performed with Trimmomatic v.0.39 (Bolger et al., 2014). Sequence assembly was performed using GetOrganelle v.1.7.7.1 (Jin et al., 2020). Alignment of these sequences together with the continental *Parnassius* sequences yielded a dataset of 15,185 bp, including indels (accession number = submitting).

### 2.4. SNP Detection

SNPs were detected from the extracted DNA using the GRAS-Di method. Enoki and Takeuchi (2018) presented the protocol at a conference, and Hosoya et al. (2019) subsequently published the detailed procedure. This protocol constructs libraries using two sequential PCR steps like MIG-seq (Suyama and Matsuki 2015). The first PCR primers consist of 10 bases of Illumina Nextera adaptor 3′-end sequences plus 3-base random oligomers (13 bases). The final PCR product is purified using columns or magnetic beads without size selection and then applied for sequencing on an Illumina platform. GRAS-Di has the advantages of simplicity in library construction and the ability to detect many SNPs. Libraries were prepared using the 2nd-step tailed PCR method; primers for 1st and 2nd PCR were followed by Hosoya et al. (2019). Cyclized DNA was prepared according to the manual using the produced library and the MGIEasy Universal Library Conversion Kit (App-A, MGI Tech). DNA Nanoballs (DNB) were prepared using the DNBSEQ DNB Rapid Make Reagent Kit (MGI Tech) and High-Throughput Pair-End Sequencing Primer Kit (App-D, MGI Tech), following the provided manuals. Sequencing was performed using DNBSEQ-T7RS High-throughput Sequencing Kit (MGI Tech) with the primer supplied with the High-Throughput Pair-End Sequencing Primer Kit (App-D) and DNBSEQ-T7 (MGI Tech) platform at a 2×150 bp condition. To remove primer sequences, the first three bases of each read were removed using Cutadapt v. 4.0 (Martin, 2011). Paired reads with a quality score of less than 30 and less than 75 bases were filtered out using Sickle v.1.33 (Joshi and Fass, 2011). Sequences after 75 nucleotides were retained to standardize read length for data analysis. The raw fastq sequence data of each sample were deposited into NCBI Sequence Read Archive (Supplemental Table 2).

Subsequently, denovo_map.pl of Stacks v. 2.62 (Catchen et al., 2013) was executed with the following parameters: minimum number of reads required to form a stack (m = 5), maximum number of nucleotide differences allowed between stacks for merging (M = 2), maximum number of nucleotide differences allowed when building the catalog (n = 2,), and assembly using paired-end reads (--paired). The SNPs information was obtained by running the program with the following settings: n = 2 (maximum number of nucleotide differences allowed when merging stacks) and assembly using paired end reads. These analyses were conducted at the commercial sequencing facility.

### 2.5. Phylogeographic Analyses

For COI and COII data, phylogenetic trees were constructed using the neighbor-joining (NJ) (Saitou & Nei, 1987), maximum likelihood (ML), and Bayesian inference (BI) methods. NJ trees were generated using MEGA X (Kumar et al. 2018) with the *p*-distance method. ML trees were constructed with RAxML (Stamatakis, 2014), and BI trees were built with MrBayes v.3.1.2 (Ronquist & Huelsenbeck, 2003). Bootstrap testing was conducted with 1,000 trials for the NJ and ML trees. ML analysis was performed using each model for each gene codon and assessed using Akaike information criterion (AIC) scores in Modeltest2 v. 2.3 (Nylander, 2004), and the selected model was HKY + I + G4. Similarly, BI was performed using each model for each gene codon and assessed using AIC scores in Kakusan4 (Tanabe, 2011), and the selected model was HKY + I + G. We aligned the sequences of 185 adults. As outgroups, we used the continental *Parnassius* sequences: *Parunassius glacialis* in China (NC 065029_1), *P. stubbendorfii* (OP 709281_1), *P. orleans* (NC 072335_1), and *P. epaphus* (NC 026864_1). Haplotype networks were obtained with TCS v. 1.21 (Clement et al., 2000).

For whole mtDNA sequence data, phylogenetic trees were constructed using the neighbor-joining (NJ) (Saitou & Nei, 1987), maximum likelihood (ML), and Bayesian inference (BI) methods. NJ trees were generated using MEGA X (Kumar et al. 2018) with the *p*-distance method. ML trees were constructed with RAxML (Stamatakis, 2014), and BI trees were built with MrBayes v.3.1.2 (Ronquist & Huelsenbeck, 2003). Bootstrap testing was conducted with 1,000 trials for the NJ and ML trees. ML analysis was performed using each model for each gene codon and assessed using Akaike information criterion (AIC) scores in Modeltest2 v. 2.3 (Nylander, 2004), and the selected model was TIM2 + G4. Similarly, BI was performed using each model for each gene codon and assessed using AIC scores in Kakusan4 (Tanabe, 2011). The selected model was GTR + G. We aligned the sequences of three adults of the three representative lineages (East Japan, West Japan, and Chugoku-Shikoku). As outgroups, we used the continental *Parnassius* sequences; *Parunassius glacialis* in China (NC 065029_1), *P. stubbendorfii* (OP 709281_1), *P. orleans* (NC 072335_1), and *P. epaphus* (NC 026864_1), *P. acdestis* (NC 072544_1), *P. hide* (NC 072287_1), *P. cephalus* (NC 026457_1), *P. simo* (NC 072286_1).

We estimated divergence times using BEAST v. 2.7.6 (Drummond et al., 2006; Drummond & Rambaut, 2007) with 10,000,000 generations of Markov-chain Monte Carlo iterations, sampled every 1,000 generations under a relaxed clock model with an uncorrelated lognormal distribution. Here, we used the HKY + G model and three calibration time points according to Si et al. (2020). The calibrations for the molecular dating were based on two butterfly fossils: *Praepapilio colorado* Durden (Papilionidae) from the Green River Shale of Colorado (USA) of mid-Eocene early Lutetian age (41– 48 MYA) and *Thaites ruminiana* Scudder (subfamily Parnassiinae) from Aix-en-Provence (southern France) of late Oligocene Chattian age (23–28 MYA) (Durden and Rose 1978, Jong 2017, Condamine et al. 2018); thus, the crown group divergence of the Parnassiinae was constrained between 23 MYA and 48 MYA. The earliest divergence time of seven *Parnassius* subgenera other than *P. epaphus,* exclusive of subgenus *Parnassius* was set to be 16–37 Ma.

### 2.6. SNP Data Analysis

We used STRUCTURE v. 2.0 (Pritchard et al., 2000) to delineate the number of genetically identified clusters (*K*) and assign individuals to clusters without prior information on their origin population. We calculated an *ad hoc* criterion of *ΔK* (Evanno et al., 2005) for determining the optimal *K* value using STRUCTURE Harvester (Earl & Holdt, 2012). The assignment index for each individual in each population was calculated using the CLUMPAK server (Kopelman et al., 2015). Genetic variation associated with various hierarchical levels of populations and their fixation index (*F*) values were quantified using analysis of molecular variance (AMOVA; Excoffier et al., 1992) in GenoDive ver. 3.0. (Meirmans, 2020). We conducted principal component analysis (PCA) using GenoDive v. 3.0.

We used Netview R v.1.0 operated by R v.4.2.3 (Neuditschko et al. 2012; Steinig et al. 2016) to reveal fine-scale population stratification independent of *a priori* ancestry information. We generated a population network based on a shared allele distance matrix (1-identity by state (IBS)) generated with PLINK v.1.9. The network was visualized by R v.4.2.3. We also used TreeMix v.1.13 (Pickrell and Pritchard, 2012) to investigate patterns of population splits and cross-population gene flow.

## 3. Results

### 3.1. Phylogeography

We detected and aligned COI (721 bps) and COII (470 bps) for each individual (accession number = in submitting). Although, if the samples in Okayama were generally included in the Chugoku Region were put into West Japan (Haplotype 06), three distinct clusters were identified in the Haplotype network by TCS (Fig. 2), corresponding to the West Japan, East Japan, Chugoku-Shikoku respectively. Demography statistics were calculated for the three strains, and the Chugoku-Shikoku strain showed a trend towards higher nucleotide diversity than the other strains (Table S1). Fu’s *F* values were significantly negative in the West Japan and East Japan lineages, while Tajima’s *D* was not significantly negative, suggesting a weak tendency towards population expansion. On the other hand, neither value was negative for the Chugoku-Shikoku lineage. Skyline plots showed recent population increases in West Japan and the Chugoku -Shikoku lineage (Fig. S1) but not in the East Japan lineage.

**Fig. 2.**
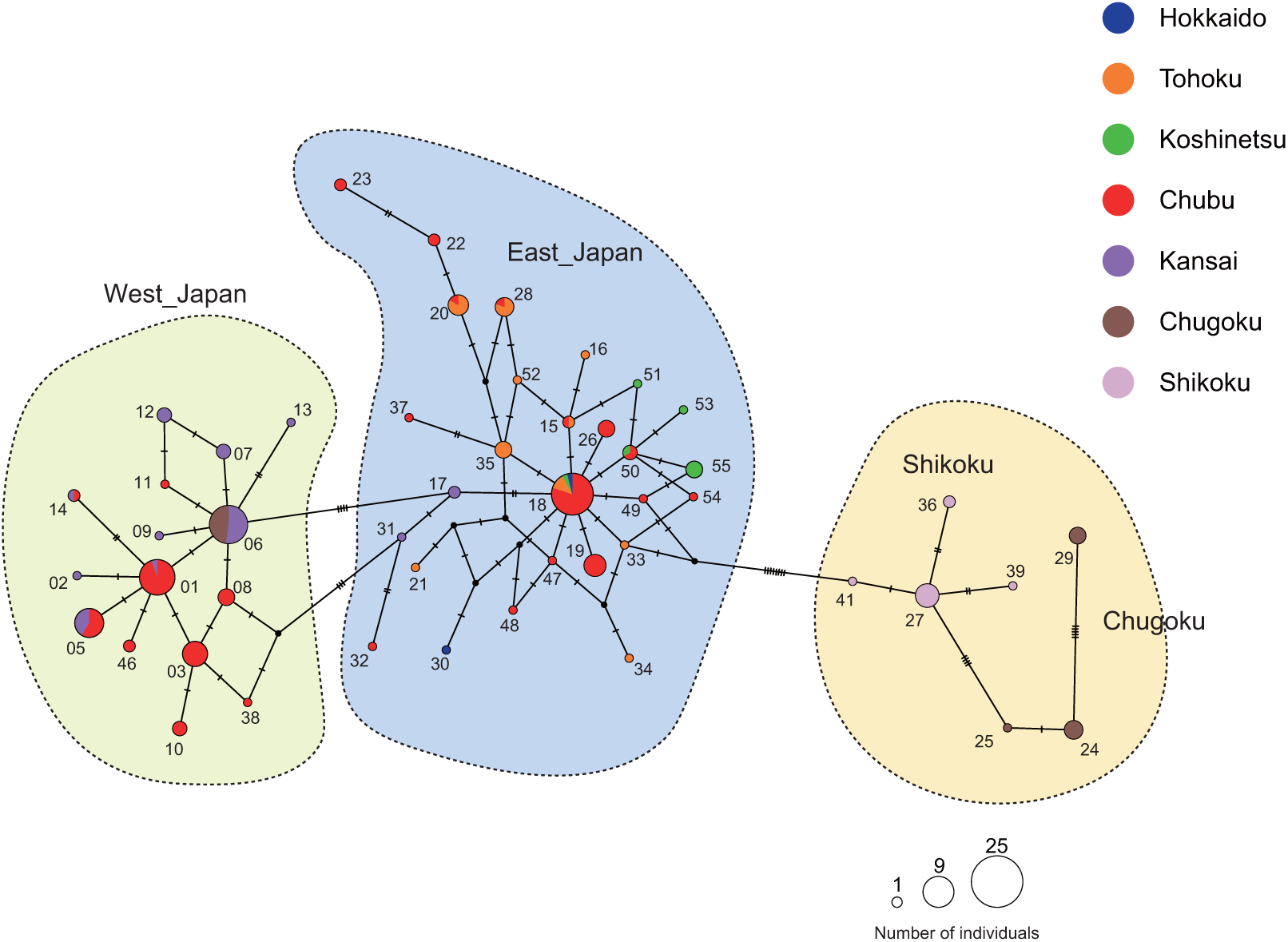
Haplotype network depicted by TCS using the sequences of 49 haplotypes detected in *Parnassius glacialis* in Japan. Each number was the haplotype number.

The three phylogenetic trees could be combined into one consensus tree (Fig. 3). There, the posterior probability and bootstrap value for the branching of the Chugoku-Shikoku lineage were highest, from which the East Japan and West Japan lineages diverged. A clear difference between the three phylogenetic trees was the position of the continental *P. glacialis*. The position of the continental *P. glacialis* was out of Japanese lineages in NJ (Fig. S2). However, the ML and BI trees contained that position within the Japanese lineages (Fig. S3 and 3).

**Fig. 3.**
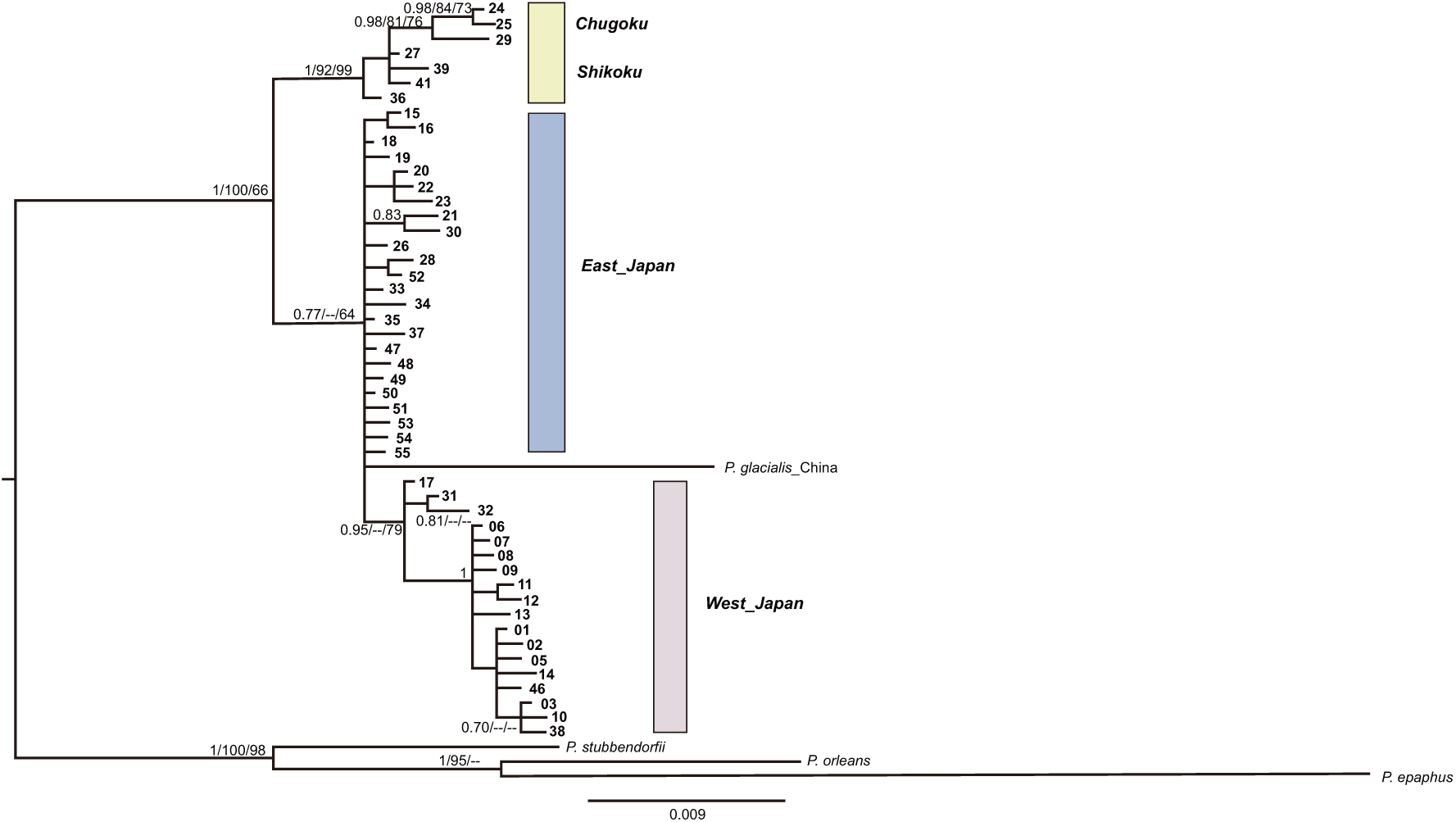
A consensus tree of BI (Bayesian Inference), ML (Maximum Likelihood), and NJ (Neighbor Joining) methods. The tree was the result of BI method. The values above each branch showed the posterior probability of BI, the bootstrap value of ML, and that of NJ, respectively. The values less than 0.6 of posterior probability and 60 of bootstrap value were not shown in this figure. The bald letters indicated the haplotype numbers.

The continental *P. glacialis* was included within *P. glacialis* in Japan, so the phylogenetic tree was rebuilt using the whole mtDNA sequences of the continental *Parnassius* species and those of three lineages of *P. glacialis* in Japan (accession number = in submitting) by the three phylogenetic methods and Beast. The consensus tree (Fig. S4) showed the continental *P. glacialis* located outside *P. glacialis* in Japan, which was derived from the continental *P. glacialis*, which was supported with high probability (1/100/100: BI/ML/NJ). The oldest lineage of *P. glacialis* in Japan was the Chugoku-Shikoku lineage (Fig. 4), which was supported with a high probability (1/100/100). *P. glacialis* in Japan diverged from the continental *P. glacialis* at 3.05 MYA ; after that, the remaining the East Japan and West Japan lineages diverged at 1.05 MYA, showing again that the Chugoku-Shikoku lineages were most ancestral ones across the Japanese Archipelago. The East Japan and West Japan lineages diverged at 0.62 MYA.

**Fig. 4.**
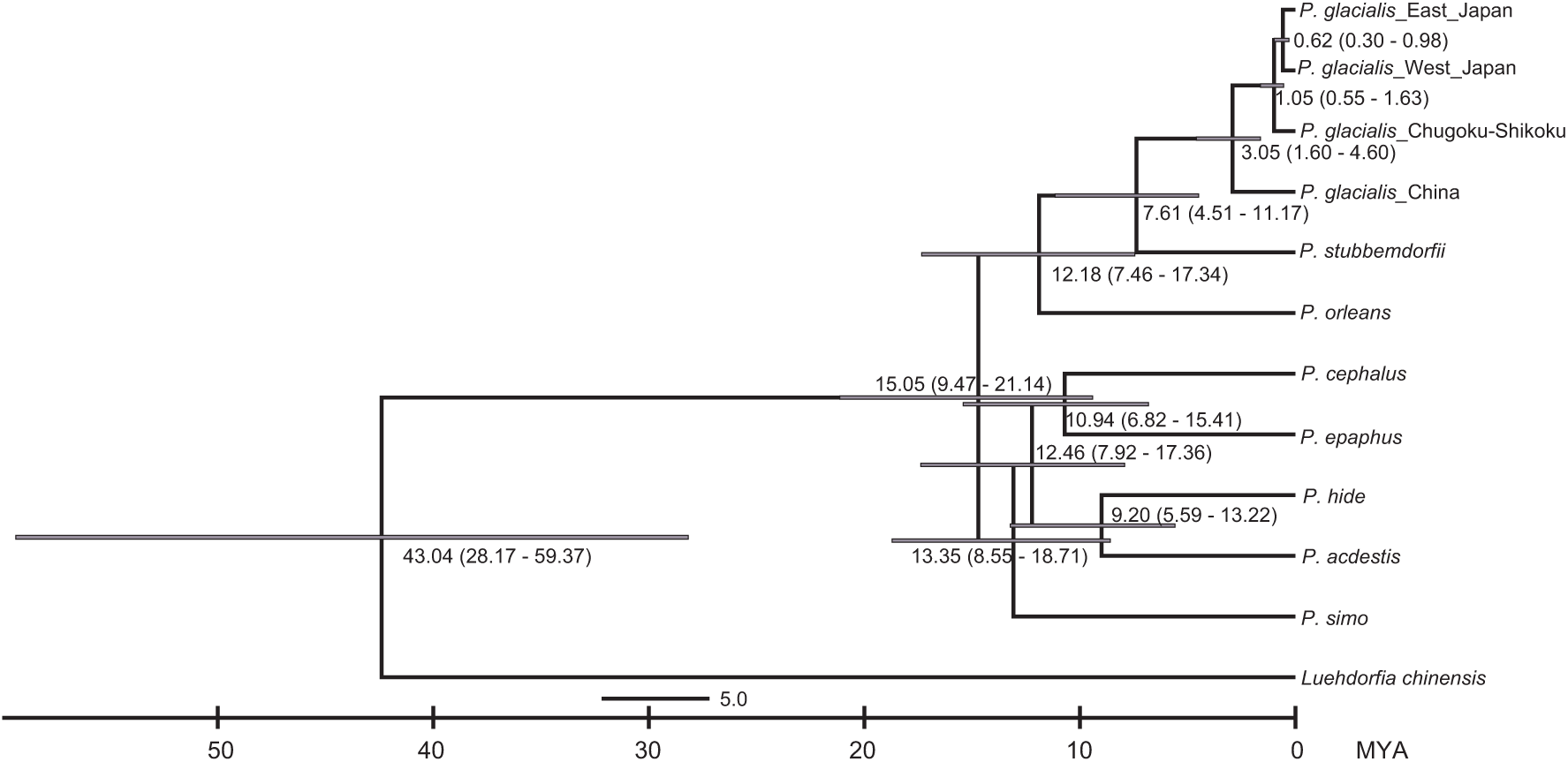
Bayesian estimates of the divergence time obtained using BEAST ver. 2.7.6. under HKY + G model, using whole mtDNA sequences of nine species and the three lineages of *P. glacialis* in Japan. The values around each node indicate the estimated divergence time. Horizontal bars indicate the 95% credible age interval at each node, with a posterior probability 1. The representative sequences for the West Japan, East Japan, and Chugoku-Shikoku lineages were obtained from the samples collected in Shiga, Hokkaido, and Kochi prefectures.

### 3.2. Population structure depicted by SNPs

We detected 3067 SNP loci using a method of GRAS-Di (Enoki and Takeuchi, 2018; Hosoya et al., 2019) and analyzed with STRUCTURE based on the information of those loci for 160 individuals. The results showed that *ΔK* (Evanno et al., 2005) for determining the optimal *K* value using STRUCTURE Harvester (Earl & Holdt, 2012) was highest at *K* = 5, which was the most appropriate cluster number (Fig. 5). Four clear genetic clusters were found at *K* = 5: Hokkaido and Tohoku (dark blue), Koshinetsu (blue), Chubu (white) and Shikoku (orange)̶in the case of *K* = 6, a new cluster appeared in Shizuoka, indicating that the populations across the Japanese Archipelago were divided into eastern and western Japan, with Shizuoka as the boundary.

**Fig. 5.**
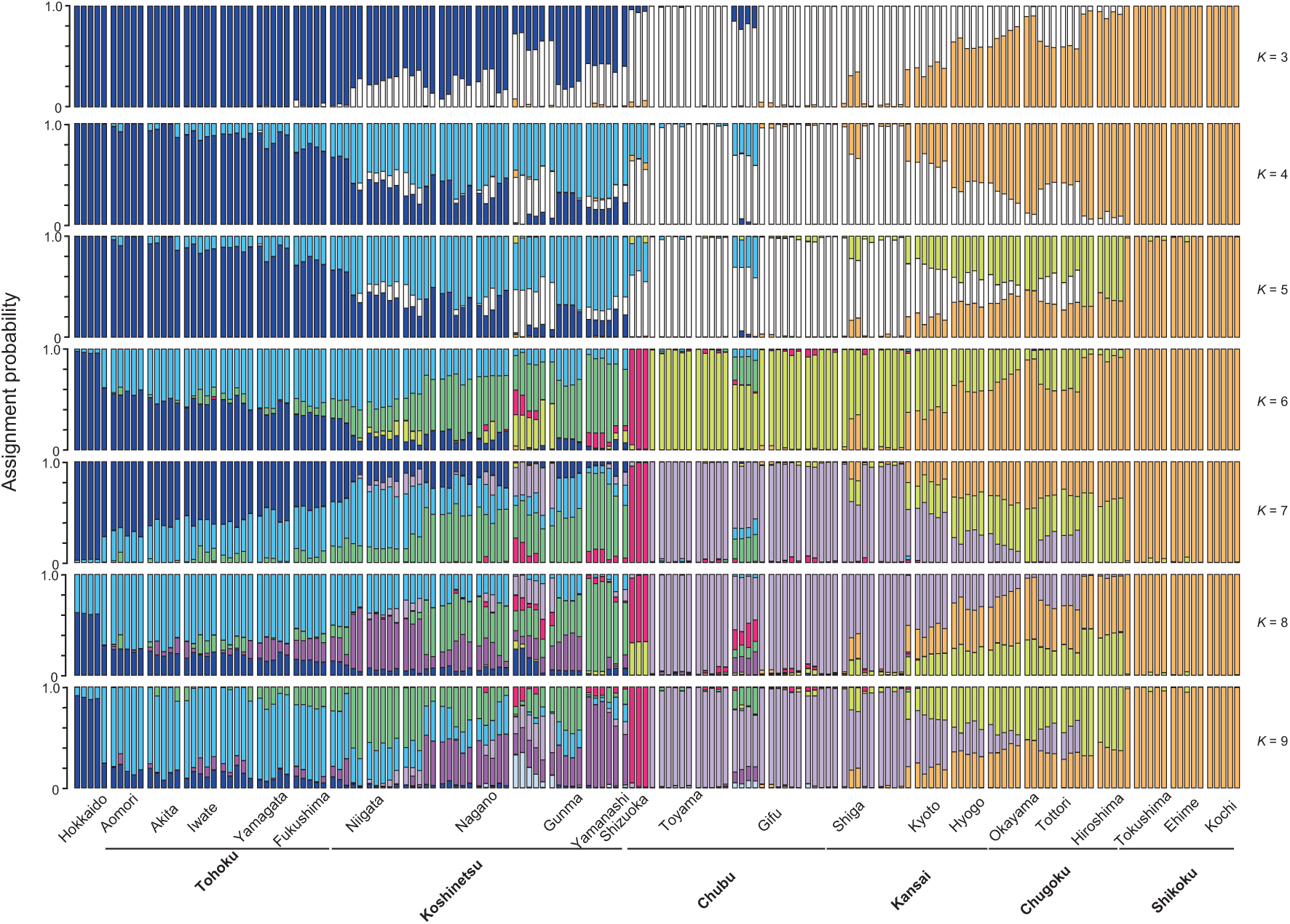
Population assignment using STRUCTURE from *K* = 3 to 9. Each population (prefecture) and regional name (in bald letters) were shown in the bottom margin from North (left) to South (right).

The results analyzed by NetviewR were consistent with those of STRUCTURE (Fig. 6). Characteristically, Hokkaido was isolated with the Blakiston Line between Hokkaido and Honshu (main Island), and the Kansai was divided into two groups. One group of the Kansai was connected to the Chugoku, while the other was connected to the Chubu. Individuals from the Kansai connected to the Chugoku were collected from the western shore side of Lake Biwa to Hyogo prefecture. In contrast, those from the Kansai connected to the Chubu were collected from the eastern shore side of Lake Biwa to Toyama prefecture, with Lake Biwa acting as one geographical barrier. Another clear barrier was found between the northern regions (Tohoku, Koshinetsu, and Chubu) and the central regions (Chubu and Kansai).

**Fig. 6.**
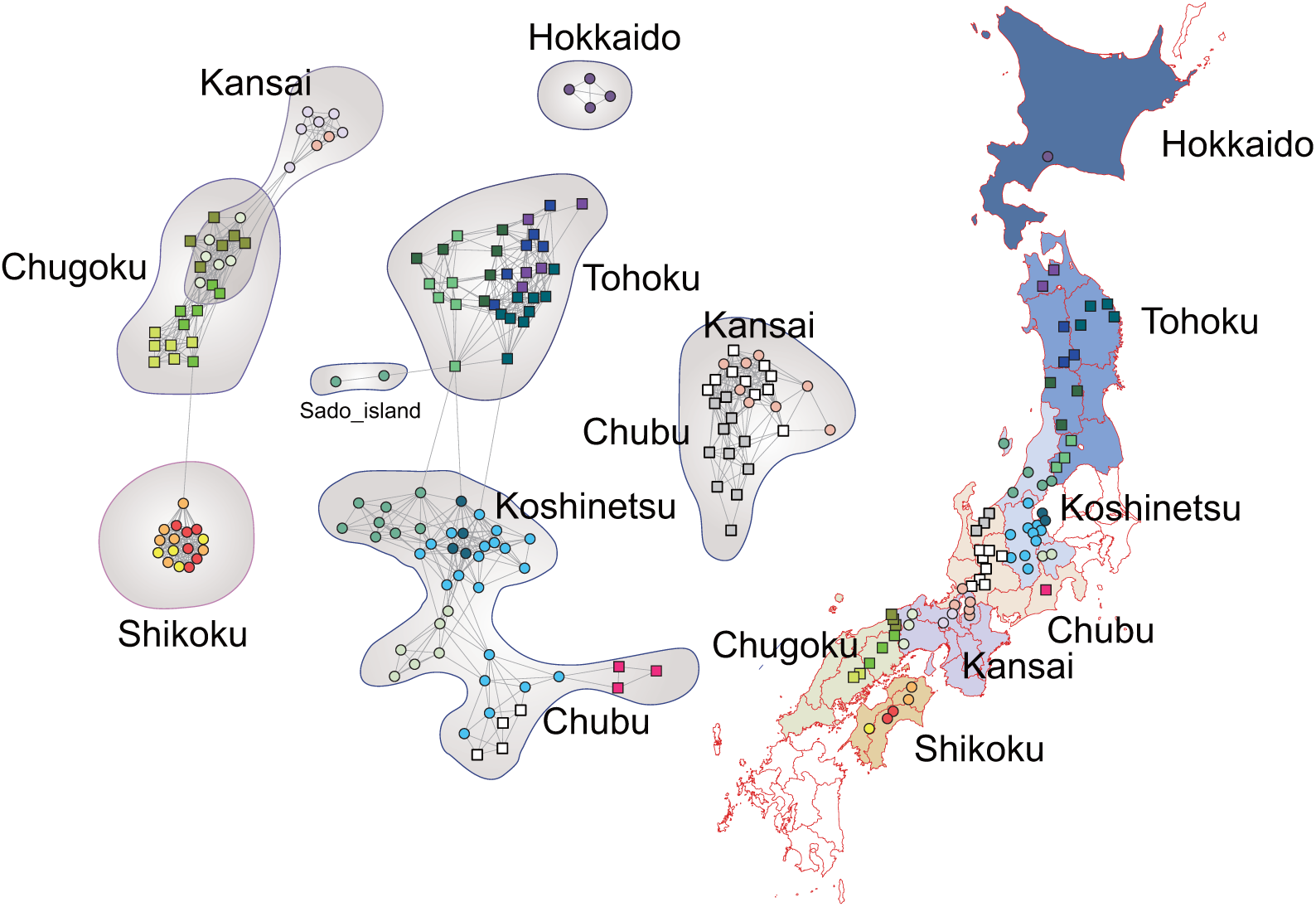
The result of NetviewR using SNPs variations. Each line showed the connection of each genotype.

The individuals from the Chubu connected to those of the Kansai, including those collected in northern Gifu and Toyama prefectures. The other individuals of the Chubu were connected to those of the Koshinetsu and Tohoku and included individuals from southern Gifu and Shizuoka. The individuals from Sado Island in the Sea of Japan were not connected to the individuals from Niigata, where it is administratively included, but rather to those from the Tohoku. These results were generally consistent with the Treemix results (Fig. S5).

The PCA results were consistent with the STRUCTURE and NetviewR results (Fig. S6). Hokkaido and Shizuoka prefectures were located separately, and the three populations in the Shikoku region also formed a single cluster.

## 4. Discussion

We revealed that *P. glacialis* in Japan was divided into three distinct genetic lineages in mtDNA of COI + COII analyses: East Japan, spanning from Hokkaido to the Koshinetsu region; West Japan, spanning Chubu to the Chugoku region; and the Chugoku-Shikoku region. The results of the SNP analyses generally supported these genetic structures. Of particular interest is that the Chugoku-Shikoku lineage was estimated to have the oldest branching time. Furthermore, the Haplotype network showed that the East Japan lineage was connected to the Chugoku-Shikoku lineage rather than the geographically closer West Japan lineage (Fig. 2).

In the phylogenetic trees with the continental *Parnassius* species as the outgroup, three clusters were generally identified in the trees from the three methods (Fig. 3). However, phylogenetic relationships other than the Chugoku-Shikoku lineage were unclear in the ML method (Fig. S3). Despite this, all phylogenetic trees suggested that the Chugoku-Shikoku lineage diverged first. The phylogenetic tree based on 1192 bp of mtDNA sometimes showed the continental *P. glacialis* included within the phylogenetic tree of *P. glacialis* in Japan, but this issue was resolved using the whole mtDNA sequence (Fig. S4). Our branching time estimates indicate that *P. glacialis* in Japan diverged from the continental *P. glacialis* approximately 3.05 MYA, with the two remaining lineages diverging at 1.05MYA from the Chugoku-Shikoku lineage, confirming that the Chugoku-Shikoku lineage is the most ancestral across the Japanese Archipelago. Furthermore, the haplotype network demonstrated that the East Japan lineage is linked to the Chugoku-Shikoku lineage rather than to the geographically closer West Japan lineage (Fig. 2). These results suggest that the East Japan lineage may have been derived from the Chugoku-Shikoku lineage, followed by the divergence of the West Japan lineage from the East Japan lineage.

Our results show that (1) the Chugoku-Shikoku lineage is the oldest lineage, and (2) the East Japan lineage is linked to the Chugoku-Shikoku lineage rather than to the geographically closer West Japan lineage. Two routes have been postulated for the expansion of biota distribution in the Japanese Archipelago: A North route and a West route (Fig. 1). Let us now consider two scenarios in which the distribution is expanded from one of these routes. For example, if we assume that the East Japan lineage expanded its distribution from the North route and then derived the West Japan lineage and the Chugoku-Shikoku lineage, the Chugoku-Shikoku lineage would be the most recent lineage. This would be inconsistent with our data, which indicates that the Chugoku-Shikoku lineage was the oldest.

Let us also consider another scenario in which the Chugoku-Shikoku lineage expanded its distribution from the West route, from which the West Japan and East Japan lineage were subsequently derived. In this scenario, the most ancestral Chugoku-Shikoku lineage would be closely related to the West Japan lineage, which would also be inconsistent with our data. In other words, assuming a single route would not resolve the discrepancy with our data. The most appropriate route to resolve these discrepancies is a scenario in which two routes have been followed. Namely, the oldest Chugoku-Shikoku lineage expanded into the Japanese Archipelago via the West route. In contrast, the East Japan lineage, which is phylogenetically most closely linked to it, expanded its distribution into the Japanese Archipelago via the North route. The scenario in which the West Japan lineage diverged from the East Japan lineage in the Japanese Archipelago is the scenario that can explain our data without contradiction. In other words, this species is thought to have differentiated from the continental *P. glacialis* in mainland China, expanded its distribution in the Japanese Archipelago via two routes, and formed a contact zone in the central mainland (Honshu).

Organisms that have differentiated traits and phylogeny from east to west within the Japanese Archipelago have been known, and it has been suggested that they may have expanded their distribution in the Japanese Archipelago via two routes, one to the North and the other to the West (Suzuki et al., 1996, 2002; Yoshimura et al., 2001; Schoville et al., 2013; Hayashi and Sota, 2014). However, neither of these phenomena has been confirmed by phylogeography. The present data are the first results supporting the two-route hypothesis.

SNP analyses with the GRAS-Di detected 3067 loci. STRUCTURE analysis indicated *K* = 5 was the most appropriate population number. Four clusters were identified at *K* = 5 (Fig. 5): Hokkaido/Tohoku (dark blue), Koshinetsu (blue), Chubu (white), and Shikoku (orange). These populations are connected from the north to the south of the Japanese Archipelago, and these boundaries may act as barriers to gene flow.

Fig. 7 compares the boundaries of the clusters as identified from mtDNA (A) and SNP (B). The mtDNA results observed two clear barriers (a and b). One barrier (b) corresponds well to the Itoigawa-Shizuoka Tectonic Line, which marks the western end of the Fossa Magna. The other barrier divides the area, including Shikoku region and Hiroshima prefectures, from the area further east. The former barrier is known to divide organisms east and west of this line in other species, and geohistorical influences have been repeatedly noted (Suzuki et al., 1996, 2002; Yoshikawa et al., 2001). Conversely, the latter barrier corresponds to the boundary between Hiroshima and Okayama Prefectures, but it has not been previously identified as a biological boundary.

**Fig. 7.**
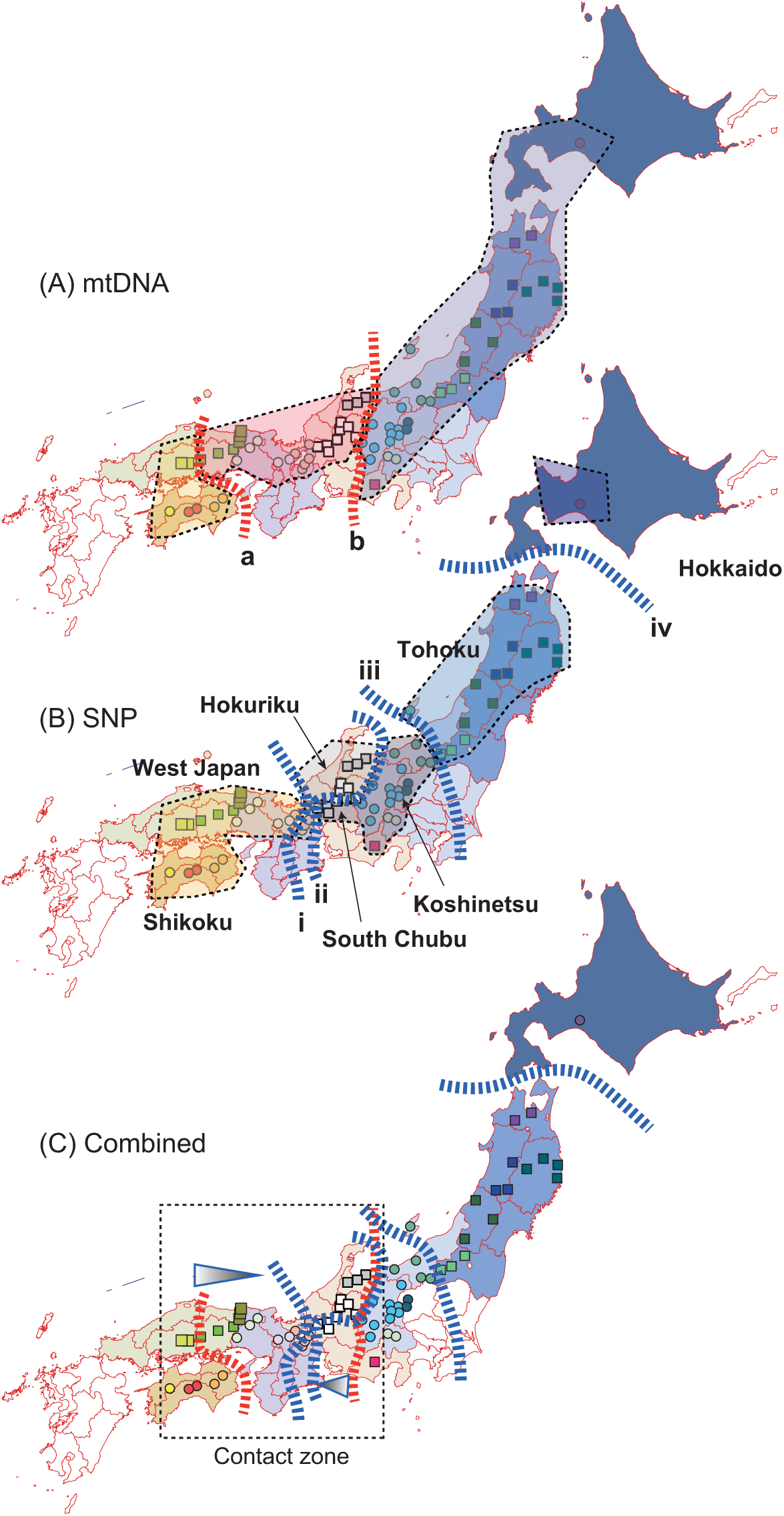
Population barriers estimated by (A) mtDNA, (B) SNP analyses. Our estimated combined barriers were shown in (C). The dotted rectangle shows inferred contact zone among the lineages.

In contrast, five genetic clusters with four barriers (i to iv in Fig. 7B) were identified in the nuclear genes of SNPs: (1) Hokkaido, (2) Tohoku, (3) East Chubu including Koshinetsu and South Chubu, (4) Hokuriku including Toyama, north Gifu and East of Shiga, (5) West Japan including West Kansai, Chugoku, and Shikoku. A closer examination reveals that barrier (i) runs through the central part of Kansai and is associated with the boundary at Lake Biwa. The areas between barriers (i) and (ii) include Toyama, northern Gifu, and the eastern shore of Lake Biwa in Shiga. Southern Gifu Prefecture, however, shows greater connectivity with the Koshinetsu region between barriers (ii) and (iii). Historically, the northern and southern administrative regions of Gifu Prefecture differed, and natural conditions remain distinct even today. This phenomenon is also observed in another butterfly, *Luehdorfia japonica*, where mtDNA haplotypes from the Chubu to the Kansai region are relatively complex and intricately intermingled in a relatively narrow range (Suzuki et al., 2023). Thus, this distribution may reflect gene flows through relatively recent human activity rather than geohistorical influences.

The combined map of barriers in Fig. 7C shows an area where the three mtDNA and SNP lineages are contacting. The area enclosed by the dotted square indicates this contact zone. In allopatric and parapatric speciation, the post-branching contact of divergent lineages is termed secondary contact (Coyne and Orr, 2004). Here, we used this term to denote the situation where different lineages of the same species are in contact. Comparing the distribution regions of mtDNA (A) and SNP (B) genotypes, SNP (B) is more finely divided. Upon closer examination, we can recognize areas of discrepancy between the two markers. For instance, the position of line (a) in mtDNA differs from that of line (i) in SNP (B). Additionally, line (b) of mtDNA is situated between lines (ii) and (iii) of SNPs. This indicates that individuals entering these two regions belong to the same group for SNPs, but to different groups for mtDNA. This indicates that SNPs are spreading across mtDNA boundaries, suggesting that the spreading nature of the gene is faster for SNPs than for mtDNA.

The triangles in Fig 7C illustrate that from the red line of the mtDNA lineage boundary, there is a nuclear gene expansion towards the blue line of the SNP boundary. This indicates that the gene flow from the boundary (a) to the east opposes the gene flow direction from the boundary (b) to the west, which corresponds to the boundary of Lake Biwa. In other words, nuclear gene flow directions appear to conflict within the contact zone. The biological significance of the Lake Biwa boundary is currently unknown but will be an area of future investigation.

A feature of this result is the difference in the degree of introgression between nuclear DNA and mtDNA: the triangles in Fig. 7C indicate regions of greater introgression of nuclear DNA than mtDNA. A factor that may contribute to this difference in the introgression of nuclear DNA and mtDNA is activity differences between males and females. The behavior of organisms can be divided into two types: male philopatry, where males tend to stay in their territories, and female philopatry, where females tend to remain in their territories (Greenwood, 1980). In the case of butterflies, including this species, most exhibit female philopatry, and the sex ratio of actively flying adults is generally skewed toward males. In this case, nuclear DNA would be expected to spread more easily.

SNP analysis indicates that the Chugoku-Shikoku lineage appears to have been extending eastward. In contrast, the East Japan lineage seems to have been expanding westward across the Sea of Japan in the past. This suggests that the two lineages currently come into contact around Lake Biwa. If this scenario holds, Haldane’s law (Coyne and Orr 2004) may be observed, where heterozygous individuals, particularly females in Lepidopteran insects, are more disadvantaged when hybridizing. However, data from amateur entomologists indicate no significant mortality increases were observed when mating individuals from distant collecting sites (Ono and Tera, personal communication). On the other hand, anecdotal evidence suggests that *P. glacialis* in Japan mates with the continental *P. glacialis* to produce viable offspring. If confirmed, this implies that reproductive isolation between the two lineages between China and Japan is not well operating and warrants careful consideration in future studies.

The present study demonstrates that *P. glacialis* colonized Japan via North and West routes to the Japanese Archipelago. While previous studies (e.g., Hayashi and Sota, 2014) inferred this dual-route entry, it has been genetically confirmed in this study. Despite undergoing long-distance migration across the Sea of Japan via these two routes, there appears to be no apparent fertility deficit in the contact zone. This study provides insights into the species’ distant history of this species. Future research on adaptive genes will be crucial for understanding their recent and future evolutionary trajectories.

## 5. Conclusion

The results of this study indicate that *Parnassius glacialis* in Japan differentiated from the continental *P. glacialis* and subsequently migrated to the Japanese Archipelago via two expansion routes: a West route and a North route. The most ancestral lineage entered the Japanese Archipelago via the West route, while its derived lineages spread via the North route. The lineage derived from the North route and the lineage from the West route are thought to be currently in contact in the central part of the Japanese Archipelago. Within the contact zone of these two lineages, differences in the distribution of nuclear genes and mtDNA were observed, suggesting that female philopatry is one of the contributing factors.

## Acknowledgments

We thank T. Takeda, N. Nagahama, T. Takeuchi, M. Kawada, H. Matsuno, A. Yokokura, T. Kudo, J. Ogasawara for their sample collection and providing samples.

## Author contributions: CrediT

H.N., A.T., K.O., T.O and K.T. conceived the study; H.T., T.N., M.H., Y.F., N.I., Y.Y., M.M., C.T., H.N., A.T., K.O., and K.T. collected the samples and extracted DNA. H.T., T.N. M.H., Y.F., N.I., Y.Y., M.M., and C.T. analysed genetic variation of mtDNA; K.Y, T.K., T.O., and K.T. conducted data analyses; K.T. wrote the manuscript; all authors approved the final version of the manuscript.

## Data accessibility

All genetic and related data used in the analyses will be deposited in Dryad.

