## supplemental_file for "Dual expansion routes likely underlie the present-day population structure in a *Parnassius* butterfly across the Japanese Archipelago": supplemental_file.pdf

Supplementary materials

Species name

*Parnassius glacialis* has been conventionally used for the Japanese clouded butterfly, but recently *P. citrinarius* has been adopted in some journals. This assumes that *P. glacialis* is a synonym of *P. citrinarius*. However, since no species description can be confirmed as the basis for this assumption, we have decided to use *P. glacialis* for convenience in this paper. The issue of this species naming should be left to taxonomists in the future.

Fig. S1. Bayesian skyline plots for the three lineages.

Fig. S2. NJ phylogenetic tree. The values above each branch showed the bootstrap value of NJ. This figure did not show the values less than 60 of bootstrap value. The bald letters indicated the haplotype numbers. *P. glacialis*_China was the sample from China.

Fig. S3. ML phylogenetic tree. The values above each branch showed the bootstrap value of ML. This figure did not show the values less than 60 of bootstrap value. The bald letters indicated the haplotype numbers. *P. glacialis*_China was the sample from China.

Fig. S4. A consensus tree of BI (Bayesian Inference), ML (Maximum Likelihood), and NJ (Neighbor Joining) methods using whole genome sequencing of mtDNA. The values above each branch showed the posterior probability of BI, the bootstrap value of ML, and that of NJ, respectively. The West Japan, East Japan, and Chugoku-Shikoku lineages were *P. glacialis*_West_Japan, *P. glacialis*_East_Japan, and *P. glacialis*_Chugoku-Shikoku, showing representative sampling regions.

Fig. S5. The result of Treemix using SNPs variations. Divergence of key populations presented as an allele-frequency-based tree with residuals of drift parameters mapped as migration edges.

Fig. S6. The result of PCA using SNPs variations.

Table S1. Genetic diversities and the results of neutrality tests of mtDNAof *Parnassius glacialis* in Japan.

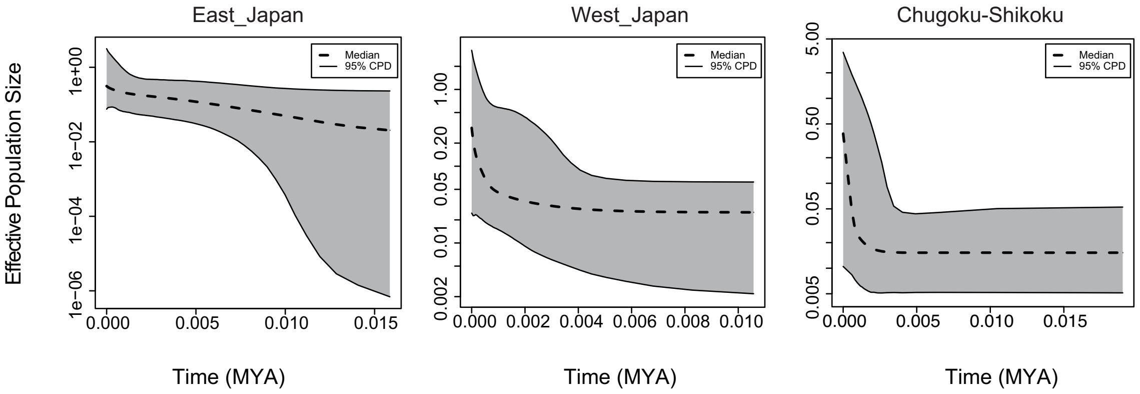
East_Japan
West_Japan
Chugoku-Shikoku
Effective Population Size
Median
95% CPD
Time (MYA)

Fig. S1

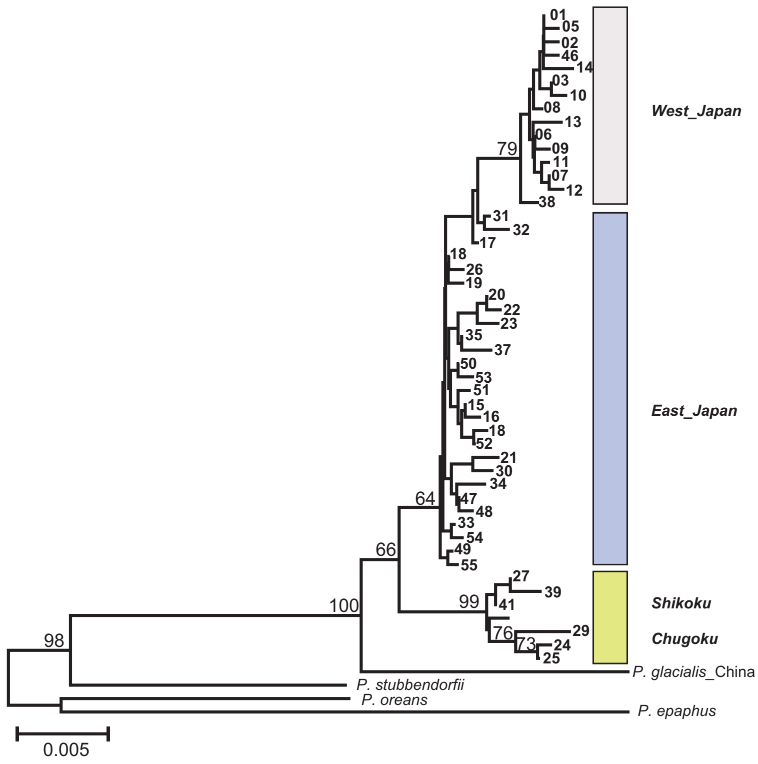
West_Japan
East_Japan
Shikoku
Chugoku
P. glacialis_China
P. stubbendorfii
P. oreans
P. epaphus
0.005

Fig. S2

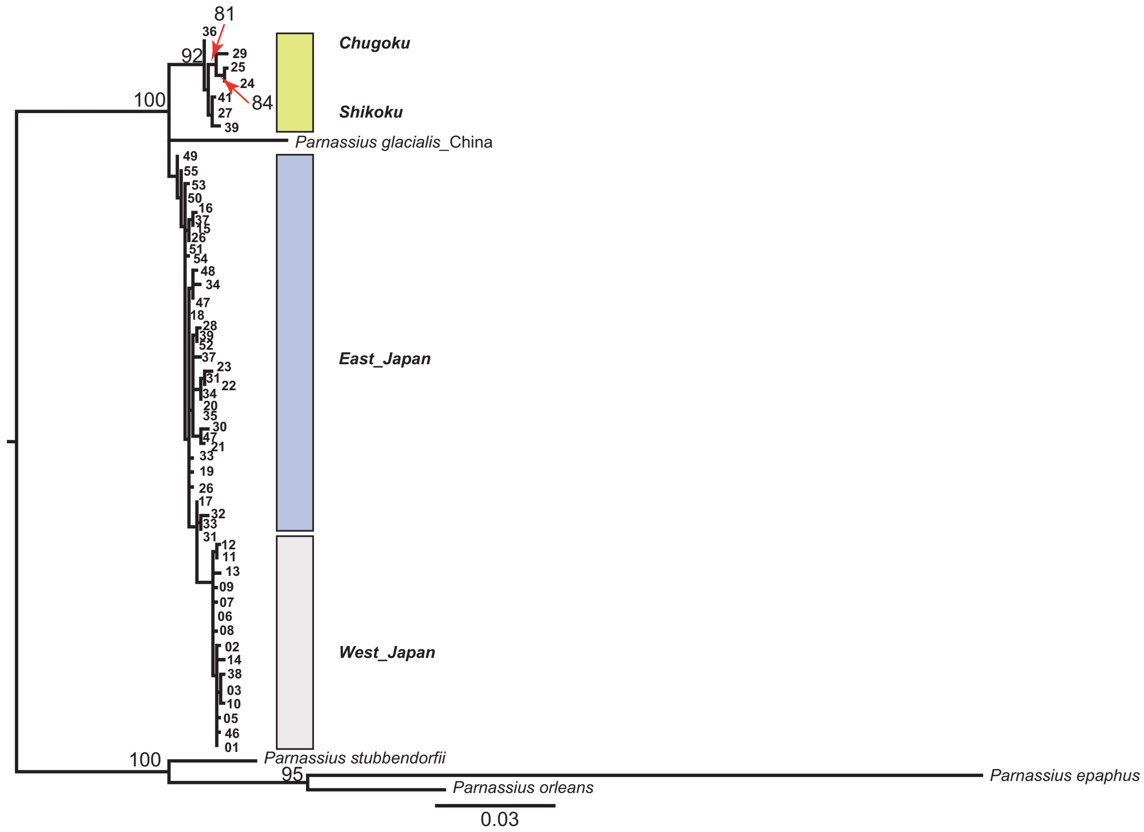
81
92
100
84
Chugoku
Shikoku
Parnassius glacialis_China
East_Japan
West_Japan
100
Parnassius stubbendorfii
95
Parnassius epaphus
Parnassius orleans
0.03

Fig. S3

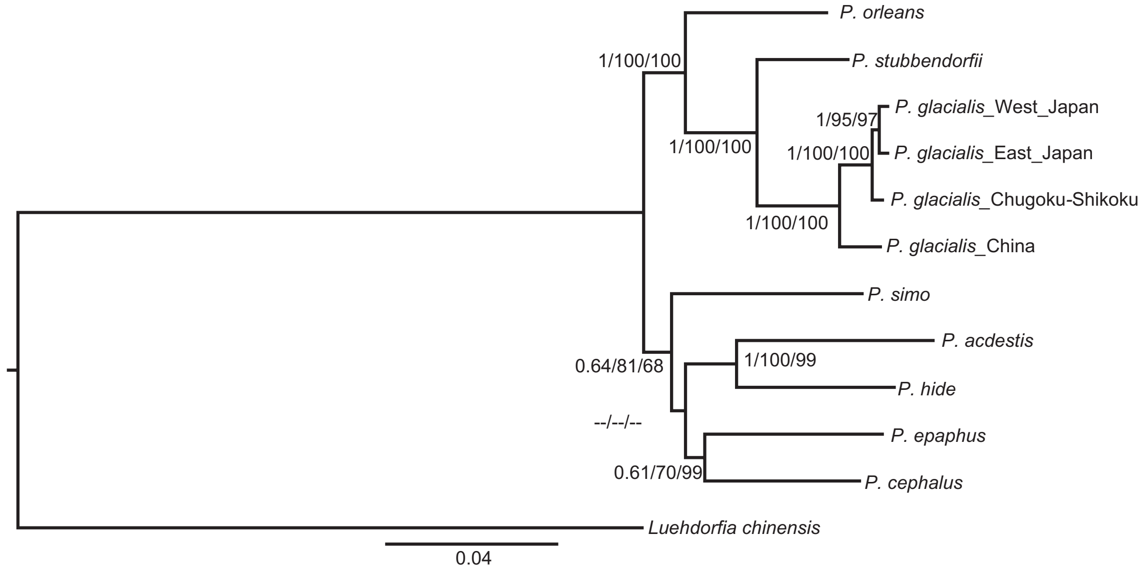
P. orleans
1/100/100
P. stubbendorfii
1/95/97
P. glacialis_West_Japan
1/100/100
1/100/100
P. glacialis_East_Japan
P. glacialis_Chugoku-Shikoku
1/100/100
P. glacialis_China
P. simo
P. acdestis
1/100/99
0.64/81/68
P. hide
--/--/--
P. epaphus
0.61/70/99
P. cephalus
Luehdorfia chinensis
0.04

Fig. S4

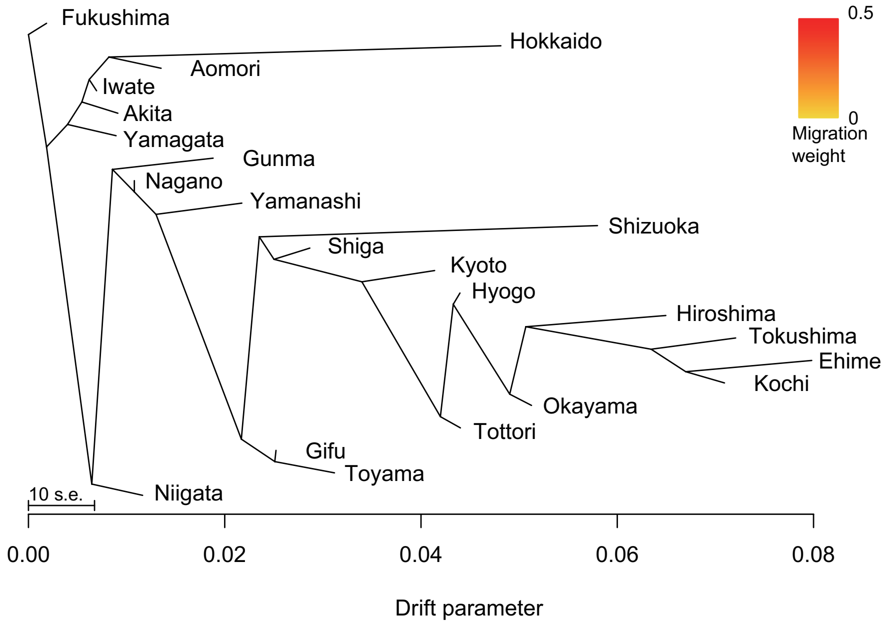
Fukushima
Hokkaido
Aomori
Iwate
Akita
Yamagata
Gunma
Nagano
Yamanashi
Shizuoka
Shiga
Kyoto
Hyogo
Hiroshima
Tokushima
Ehime
Kochi
Okayama
Tottori
Gifu
Toyama
Niigata
10 s.e.
0.00
0.02
0.04
0.06
0.08
Drift parameter
0.5
0
Migration weight

Fig. S5

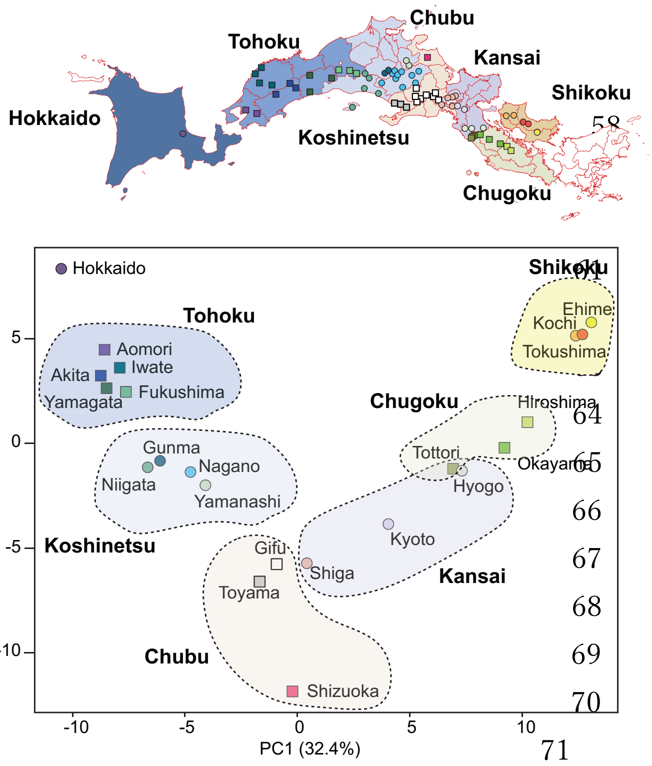
Tohoku
Chubu
Kansai
Shikoku
Hokkaido
Koshinetsu
Chugoku
Hokkaido
Tohoku
Aomori
Iwate
Akita
Fukushima
Yamagata
Shikoku
Ehime
Kochi
Tokushima
Chugoku
Hiroshima
Tottori
Okayama
Hyogo
Gunma
Nagano
Niigata
Yamanashi
Koshinetsu
Kyoto
Gifu
Shiga
Toyama
Kansai
Chubu
Shizuoka
PC2 (11.6%)
PC1 (32.4%)
5
0
-5
-10
-10
-5
0
5
10

Fig. S6

Table S1. Genetic diversities and the results of neutrality tests of mtDNAof *Parnassius glacialis* in Japan.

| | No. of individuals | No. of haplotypes | Nucleotide diversity ($\pi$) | Haplotype diversity (*Hd*) | Fu's *F* | *P* | Tajima's *D* | *P* |
|---|---|---|---|---|---|---|---|---|
| East_Japan | 81 | 27 | 0.0021 | 0.89 | -19.99 | 0 | -1.11 | 0.13 |
| West_Japan | 82 | 14 | 0.0014 | 0.86 | -6.78 | 0.006 | -1.17 | 0.11 |
| Chugoku-Shikoku | 22 | 7 | 0.0032 | 0.81 | 0.95 | 0.7 | 0.51 | 0.73 |
